# Pollutants and insecticides drive local adaptation in African malaria mosquitoes

**DOI:** 10.1101/067660

**Authors:** Colince Kamdem, Caroline Fouet, Stephanie Gamez, Bradley J. White

## Abstract

The *Anopheles gambiae* complex contains a number of highly anthropophilic mosquito species that have acquired exceptional ability to thrive in complex human habitats. Thus, examining the evolutionary history of this Afrotropical mosquito may yield vital information on the selective processes that occurred during the adaptation to human-dominated environments. We performed reduced representation sequencing on 941 mosquitoes of the *Anopheles gambiae* complex collected across four ecogeographic zones in Cameroon. We find evidence for genetic and geographic subdivision within *An. coluzzii* and *An. gambiae* sensu stricto – the two most significant malaria vectors in the region. Importantly, in both species, rural and urban populations are genetically differentiated. Genome scans reveal pervasive signatures of selection centered on genes involved in xenobiotic resistance. Notably, a selective sweep containing detoxification enzymes is prominent in urban mosquitoes that exploit polluted breeding sites. Overall, our study suggests that recent anthropogenic environmental modifications and widespread use of insecticides are driving population differentiation and local adaptation in vectors with potentially significant consequences for malaria epidemiology.

## INTRODUCTION

Humans and their actions on ecosystems can profoundly alter the evolutionary trajectory of other organisms in various ways (Palumbi 2001; Bull and Maron 2016; Hendry et al. 2017). Signatures of human-driven evolutionary changes are particularly evident in cities and urbanizing regions, which often necessitate a rapid adaptation to complex environments (Alberti 2015; Alberti et al. 2017; Hendry et al. 2017). A wealth of data exists on genetic and phenotypic signals associated with adaptation to anthropogenic changes in multiple taxa, but the extent of signatures across genomes remain largely unexplored. The widespread use of xenobiotics in agriculture and public health also increases selective pressures on pathogens and their vectors, prompting significant evolutionary adjustments that have yet to be fully elucidated (Georghiou 1972; Mallet 1989; Palumbi 2001). The application of powerful genomic tools now available will help identify the genes and genomic regions underlying adaptation to human-induced selective pressures.

The most effective malaria mosquitoes of the Afrotropical region are remarkably anthropophilic and closely associated with human habitats (White 1974; Coluzzi et al. 1979; Antonio-Nkondjio et al. 2006; Sinka et al. 2010; Dia et al. 2013). Therefore, dissecting the genome of contemporary populations of these species may provide significant insights into the genetic targets of recent selection driven by anthropogenic disturbance. Adaptation to human-dominated environments is challenging and very selective even among highly specialized mosquitoes that feed almost exclusively on human hosts. For example, tens of *Anopheles* species transmit human malaria parasites across the continent, but only four members of the *Anopheles gambiae* complex thrive in dense urban areas (Antonio-Nkondjio et al. 2005; Oyewole and Awolola 2006; Mourou et al. 2010; Dabiré et al. 2012; Kamdem et al. 2012). The recent scaling up of measures using insecticides and insecticide-treated materials to combat malaria expose mosquitoes to an even more disruptive adaptive challenge (Sokhna et al. 2013; WHO 2013). Although the genetic basis remains unknown, studies suggest that both physiological and ecological limits and species ranges are profoundly altered in *Anopheles* mosquitoes as a result of rapid urbanization and widespread use of insecticides (Jones et al. 2012; Kamdem et al. 2012; Mwangangi et al. 2013; Tene Fossog et al. 2013; Antonio-Nkondjio et al. 2015).

The *Anopheles gambiae* complex is a group of at least nine isomorphic mosquito species exhibiting varying degrees of geographic and reproductive isolation (Riehle et al. 2011; Coetzee et al. 2013; Fontaine et al. 2015). Owing to ongoing adaptive speciation, the principal vectors of the complex have an amazing plasticity in adjusting to diverse ecological conditions, allowing them to track humans across most of tropical Africa (Davidson 1964; Coluzzi et al. 1979; Coluzzi et al. 2002). This adaptive flexibility and the underlying behavioral complexity have undermined at least one major effort at vector control (Molineaux and Gramiccia 1980). Importantly, the epidemiological consequences of adaptive radiation will be even more challenging to tackle if a part of the divergence recently described within the particularly efficient vectors *An. gambiae* sensu stricto (hereafter *An. gambiae*) and *An. coluzzii* is driven by human-induced changes (Touré et al. 1998; Wondji et al. 2005; Slotman et al. 2007; Pinto et al. 2013; Caputo et al. 2014). This hypothesis has been formulated based on ecological observations and population structure, but the genetic underpinnings still need to be clearly assessed by examining patterns of variation across the genome (Coluzzi et al. 2002; Kamdem et al. 2012; Caputo et al. 2014).

To delineate the population structure and the role of recent processes – mainly urbanization and the rapid scaling up of insecticide-treated bed net coverage – in the genetic differentiation, we genotyped 941 mosquitoes of the *An. gambiae* complex collected from diverse environments in Cameroon at >8,000 SNP. We found strong evidence for complex geographic and genetic structuring giving rise to seven subpopulations distributed along a continuum of genomic differentiation within both *An. gambiae* and *An. coluzzii*. By combining genomic evidence and ecological knowledge, we revealed that the strongest correlates of genetic differentiation in this region are intense suburban agriculture, urbanization, insecticides and forest-savannah subdivisions. These findings add to a growing body of data indicating that the inevitable rise of insecticide resistance together with ongoing local adaptation of *Anopheles* species in urban areas are the major challenges for current vector control strategies.

## RESULTS

### *Population structure of* An. gambiae *sensu lato (s.l.) sibling species*

We performed extensive sampling of human-associated *Anopheles* across the main ecological zones in Cameroon (Table S1, Figure S1) to collect diverse populations belonging to the four species of the *An. gambiae* complex that are known to occur in the country (Simard et al. 2009). Certain subpopulations or species can be overlooked when sampling is biased toward one type of population (Riehle et al. 2011). To maximize the chances that our samples best represent the genetic diversity within each species, we used several sampling methods (Service 1993) to collect both larvae and adult populations. In addition, populations of *An. gambiae* and *An. coluzzii* segregate along urbanization gradients, which seem to be the most important driver of ecological divergence in the forest zone (Kamdem et al. 2012). To validate this hypothesis and to investigate the genomic targets of local adaptation in urban environments, we surveyed several neighborhoods representing the urban and suburban ecosystems in the two biggest cities of the forest area: Douala and Yaoundé (Figure S1).

To assess the genetic relatedness among individuals, we subjected all 941 mosquitoes that were morphologically identified as *An. gambiae* s.l. to population genomic analysis. Individual mosquitoes were genotyped in parallel at a dense panel of markers using double-digest restriction associated DNA sequencing (ddRADseq), which enriches for a representative and reproducible fraction of the genome that can be sequenced on the Illumina platform (Peterson et al. 2012). After aligning ddRADseq reads to the *An. gambiae* reference genome, we used STACKS (Catchen et al. 2011; Catchen et al. 2013) to identify 8,476 SNPs (~1 SNP every 30kb across the genome) within consensus RAD loci present in more than 80% of individuals, and we inferred the population structure based on these variations. First, we performed principal component analysis (PCA) across all 941 individuals (Figure 1A). The top three components explain 28.4% of the total variance and group individuals into five main clusters. Likewise, a neighbor-joining (NJ) tree, based on Euclidian distance of allele frequencies, shows five distinct clades of mosquitoes (Figure 1B). We hypothesized that these groups at least partially correspond to the four sibling species – *An. gambiae, An. coluzzii, An. arabiensis,* and *An. melas* – known to occur in Cameroon. To confirm, we typed a subset of 288 specimens using species identification PCRs (Fanello et al. 2002; Santolamazza et al. 2004) and found that each cluster comprised a single species. In agreement with previous surveys (Wondji et al. 2005; Simard et al. 2009), our collections indicate that the brackish water breeding *An. melas* is limited to coastal regions, while the arid-adapted *An. arabiensis* is restricted to the savannah. In contrast, *An. gambiae* and *An. coluzzii* are distributed across the four eco-geographic zones of Cameroon (Figure 1D). Lee et al. 2013 recently reported frequent bouts of hybridization between *An. gambiae* and *An. coluzzii* in Cameroon. While both the PCA and NJ trees clearly separate the two species, the PCA does show intermixing of some rare individuals consistent with semi-permeable species boundaries.

**Figure 1.**
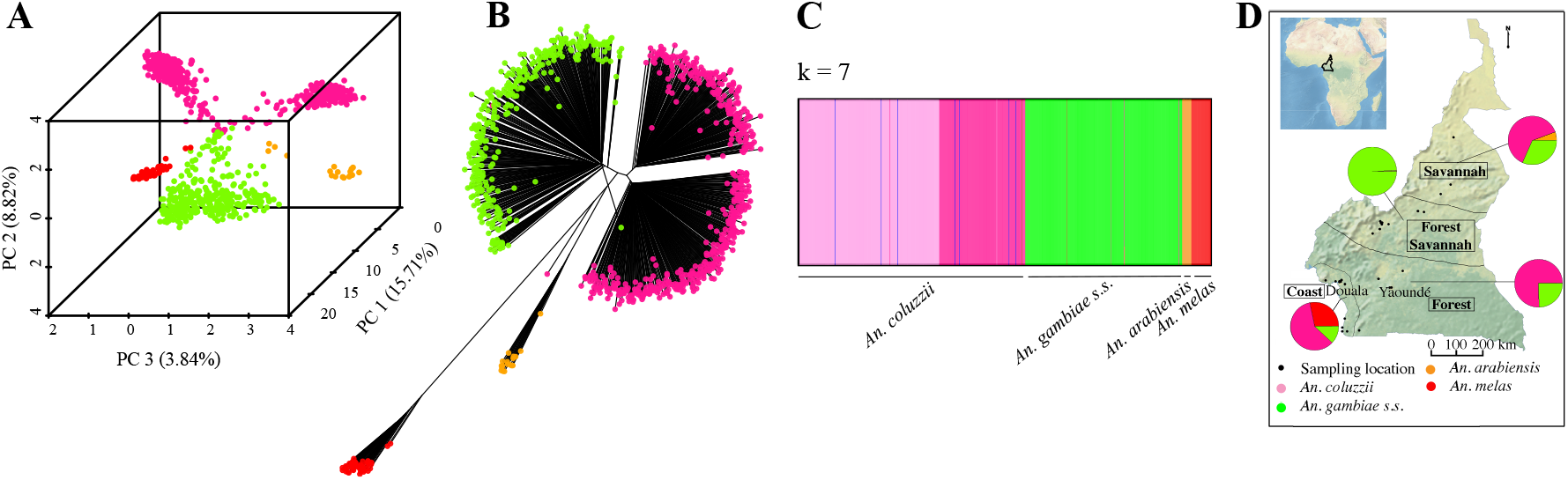
*Anopheles gambiae* complex sibling species are genetically distinct. A) PCA B) NJ Tree, and C) fastSTRUCTURE analyses clearly separate the four *An. gambiae* complex species that occur in Cameroon into discrete genetic clusters. Additional subdivision below the species level is apparent within *An. coluzzii* and *An. gambiae*. D) Species composition varies strongly between eco-geographic regions. Sampling sites are denoted by black dots.

In support of population structuring below the species level, Bayesian clustering analysis with fastSTRUCTURE (Raj et al. 2014) finds that 7 population clusters (*k*) best explain the genetic variance present in our data (Figure 1C, Figure S2). Indeed, grouping of samples within *An. gambiae* and *An. coluzzii* clades suggests that additional subdivision may exist within each species (Figure 1A, 1B). Ancestry plots further support inference from the PCA and NJ tree: at least two subgroups compose *An. coluzzii* and *An. gambiae* individuals show admixed ancestry, while *An. arabiensis* and *An. melas* are panmictic (Figure 1C, Figure S2). A cryptic subgroup of *An. gambiae* has been discovered recently by comparing indoor and outdoor fauna from the same village in Burkina Faso (Riehle et al. 2011). Visual inspections of our PCA, NJ and fastSTRUCTURE clustering results do not indicate any genetic subdivision based on the collection methods or the developmental stage. To explicitly test for the effects of the sampling methods and the geographic origin of samples on the genetic variance among individuals, we applied a hierarchical Analysis of Molecular Variance (AMOVA) (Excoffier et al. 1992), which confirmed the absence of genetic structuring based on microhabitats or temporal segregations in both species. The large majority of the genetic variation is attributable to differences among individuals. While the geographic origin accounts for 13.2 % (*p* < 0.001) and 9.4% (*p* < 0.001) of the variance in *An. coluzzii* and *An. gambiae* respectively, the effect of the types of sample is marginal and not significant.

### *Population Structure within* An. gambiae

To further resolve the population structure within 357 *An. gambiae* specimens, we performed population genetic analysis with a set of 9,345 filtered SNPs. Using a combination of PCA, NJ trees, and ancestry assignment, we consistently identify three distinct subgroups within *An. gambiae* (Figure 2, Figure S2). The first and largest subgroup (termed *GAM1*) comprises the vast majority of all *An. gambiae* specimens including individuals collected in all four eco-geographic regions (Table S1). A total of 17 individuals make up a second small subgroup (termed *GAM2*). Interestingly, individuals assigned to this cluster include both larvae and adults collected in 3 different villages spread across 2 eco-geographic regions. In the absence of any obvious evidence of niche differentiation between *GAM1* and *GAM2*, it is unclear what is driving and/or maintaining divergence between the two sympatric subgroups. Specimens collected from Nkolondom, a suburban neighborhood of Yaoundé where larval sites associated with small-scale agricultural irrigation are common (Fossog Tene et al. 2013; Nwane et al. 2013), form a genetically distinct third subgroup (termed *Nkolondom*) that appears to be a locally-adapted ecotype.

**Figure 2.**
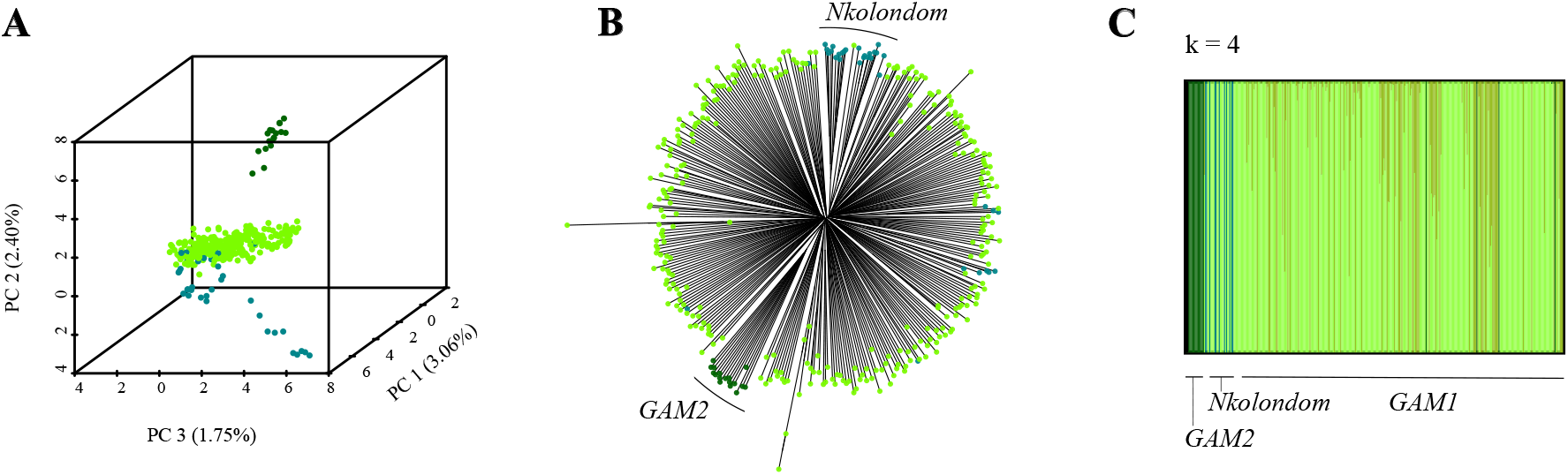
*Anopheles gambiae* is divided into three subpopulations. A) PCA, B) NJ Tree, and C) fastSTRUCTURE analyses reveal subdivisions within *An. gambiae*. We term the most abundant group *GAM1*, while a second small, but widely distributed group, is termed *GAM2*. Finally, most individuals from the village of Nkolondom are genetically distinct from other *An. gambiae* suggestive of local adaptation.

### Population Structure within *An. coluzzii*

To examine population structure within 521 *An. coluzzii* specimens, we utilized 9,822 SNPs that passed stringent filtration. All analyses show a clear split between individuals from the northern savannah region and the southern three forested regions of Cameroon (Coastal, Forest, Forest-Savannah) (Figure 3A-C). In principle, the north-south structuring could be caused solely by differences in chromosome 2 inversion frequencies, which form a cline from near absence in the humid south to fixation in the arid north (Simard et al. 2009; Fouet et al. 2012). However, we find SNPs from all five chromosomal arms consistently separate northern and southern mosquitoes, indicating a substantial genome-wide divergence between the two populations (Figure S3).

**Figure 3.**
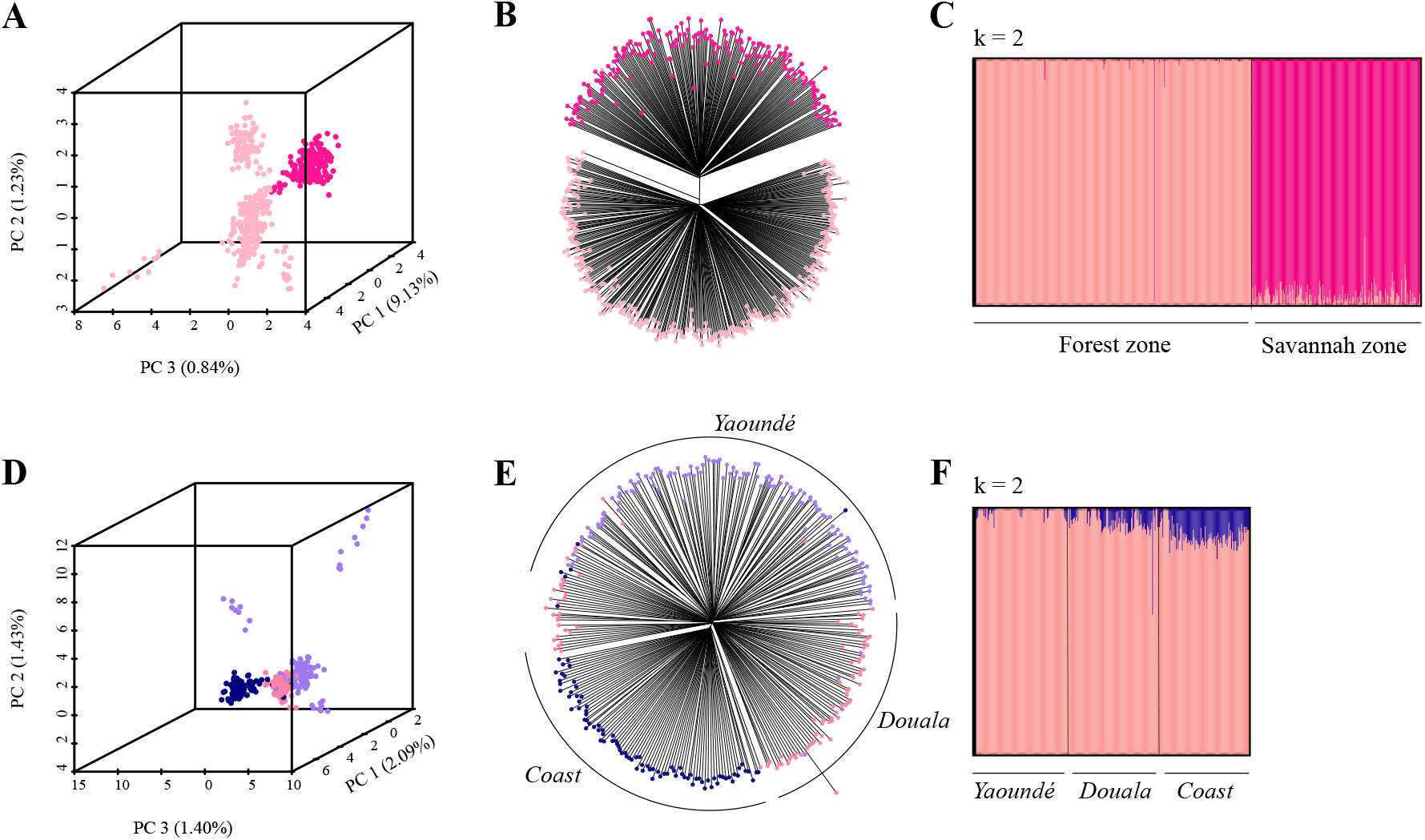
*Anopheles coluzzii* is divided into four subgroups. A) PCA, NJ Tree, and C) fastSTRUCTURE reveal major population structuring between *An. coluzzii* from the northern Savannah eco-geographic region and *An. coluzzii* from the southern three forested regions. Within the south, D) PCA, E) NJ Tree, and F) fastSTRUCTURE analyses separate mosquitoes based on geographic origin, although clustering is not fully discrete indicating a dynamic interplay between local adaptation and migration.

Southern populations of *An. coluzzii* were collected from three different areas: Douala (the largest city of Cameroon), Yaoundé (the second largest city) and the rural coastal region. PCA, NJ trees, and fastSTRUCTURE show clear clustering of southern samples by collection site (Figure 3D-F). Mosquitoes from Douala, situated on the coastal border, contain a mixture of urban and coastal polymorphisms as illustrated by their intermediate position along PC3 (Figure 3D). Despite considerable geographic segregation, clusters are not fully discrete, likely owing to substantial migration between the three sites. Taken together, the data suggest a dynamic and ongoing process of local adaptation within southern *An. coluzzii*. In contrast, no similar geographic clustering is observed in northern populations (Figure S2, S4). Slight variations can be observed between genetic clustering patterns suggested by different methods. We have only considered subdivisions that were consistent across the three methods we employed. For example, individuals from Yaoundé have been treated as a single subgroup, although PCA showed that a few specimens were relatively detached from the main cluster. This putative subdivision was not supported by the NJ tree and fastSTRUCTURE analyses (Figure 3D-F). All populations described as “urban” were collected from the most urbanized areas of the city, which are characterized by a high frequency of built environments as described in Kamdem *et al*. 2012.

### Relationships Between Species and Subgroups

Population genomic analysis identified four different *An. gambiae* s.l. species present within our samples. Within *An. gambiae* and *An. coluzzii* we identified seven potential subgroups with apparently varying levels of divergence. To further explore the relationships between different populations, we built an unrooted NJ tree using pairwise levels of genomic divergence (*F_ST_*) between all species and subgroups (Figure 4, Table S2). As previously observed in a phylogeny based on whole genome sequencing (Fontaine et al. 2015), we find that *An. melas* is highly divergent from all other species (*F_ST_* ~ 0.8), while *An. arabiensis* shows intermediate levels of divergence (*F_ST_* ~ 0.4) from *An. gambiae* and *An. coluzzii*. As expected, the sister species *An. gambiae* and *An. coluzzii* are more closely related to each other (*F_ST_* ~ 0.2) than any other species. When examining differentiation between subgroups within *An. coluzzii*, we find that the southern and northern subgroups are relatively divergent (*F_ST_* > 0.1), while differentiation between local ecotypes within the south is much lower (*F_ST_* < 0.04), corroborating patterns of population structure described within this species (Figure 3D-F). Similarly, as suggested by the genetic structuring, the level of differentiation between *An. gambiae* subgroups *GAM1* and *GAM2* is relatively significant (*F_ST_* ~ 0.1), while the suburban ecotype from Nkolondom shows a low level of divergence from *GAM1 (F_ST_* ~ 0.05). To further examine the degree of isolation of subgroups within species, we assessed the reductions in observed heterozygosity with respect to that expected under Hardy–Weinberg Equilibrium among *An. coluzzii* and *An. gambiae* populations, by computing the average Wright’s inbreeding coefficient, *F_IS_*, across genome-wide SNPs. Values of *F_IS_* close to 1 indicate a deviation from Hardy-Weinberg Equilibrium and the existence of cryptic subdivisions while *F_IS_* close to 0 suggest that there are no barriers to gene flow. In spite of the strong population genetic structure observed within *An. coluzzii* and *An. gambiae*, we found surprisingly low genome-wide *F_IS_* values (less than 0.0003) in both species, suggesting a lack of assortative mating. Overall, there is no evidence for reproductive isolation within *An. gambiae* and *An. coluzzii* despite ongoing local adaptation and significant population differentiation.

**Figure 4.**
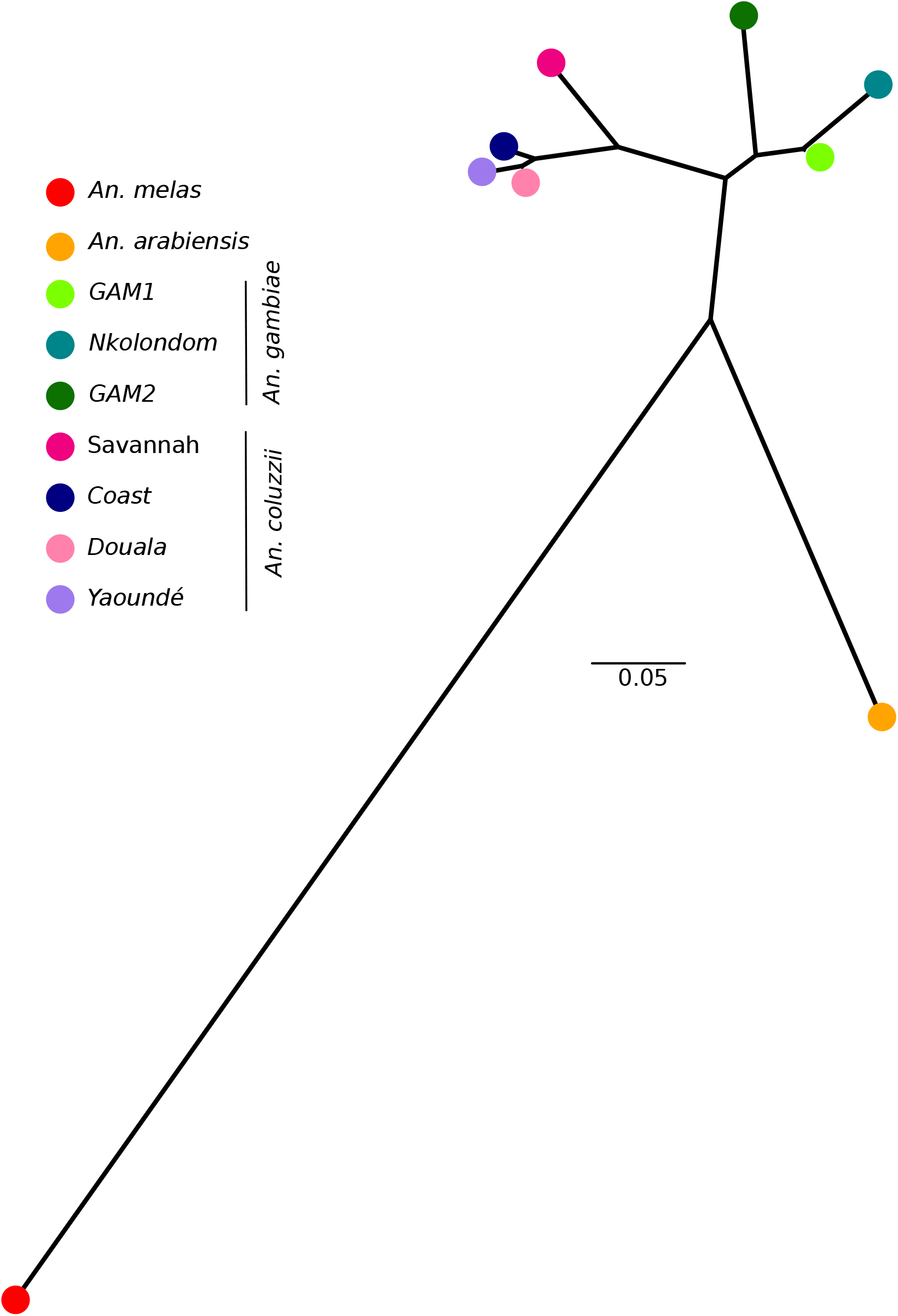
The genetic distance (*F_ST_*) between populations show recent radiation within *An. gambiae* and *An. coluzzii* clades. In an unrooted, *F_ST_*- based NJ tree, *An. melas* is most distant from all other species, while *An. gambiae* and *An. coluzzii* are sister species. Southern populations of *An. coluzzii* are more closely related to each other than to the northern savannah population. In contrast to geographic distance, the Douala subpopulation is genetically closer to Yaoundé rather than Coastal mosquitoes. Within *An. gambiae*, a relatively deep split is present between *GAM2* and *GAM1*, while *Nkolondom* appears to have recently diverged from *GAM1*.

### Signatures of selection

To find potential targets of selection within subgroups, we performed scans of nucleotide diversity (*θ_w_, θ_π_*) and allele frequency spectrum (*Tajima’s D*) in non-overlapping 150-kb windows across the genome. Scans of *θ_W_, θ_π_, and Tajima’s* D were conducted by importing aligned, but otherwise unfiltered, reads directly into ANGSD, which uses genotype likelihoods to calculate summary statistics (Korneliussen et al. 2014). Natural selection can increase the frequency of an adaptive variant within a population, leading to localized reductions in genetic diversity as the haplotype containing the adaptive variant(s) sweeps towards fixation (Maynard Smith and Haigh 1974; Tajima 1989). Selective processes can also promote the coexistence of multiple alleles in the gene pool of a population (balancing selection). Thus, genomic regions harboring targets of selection should exhibit extreme values of diversity and allele frequency spectra relative to genome-wide averages (Storz 2005).

We also performed genome scans using both a relative (*F_ST_*) and absolute (*d_xy_*) measure of divergence calculated with STACKS and *ngsTools* (Fumagalli et al. 2014), respectively. For both diversity and divergence scans we used a maximum of 40 mosquitoes per population, prioritizing individuals with the highest coverage in populations when sample size exceeded 40. In contrast to Tajima’s *D* and *F_ST_*, the genome-wide distribution of *d_xy_* and nucleotide diversity in 150-kb sliding windows yielded relatively noisy patterns due to a large variance (Wakeley 1996) (Figure 5 and 6). As a result, the identification of signatures of selection was based primarily on outliers of Tajima’s *D* and *F_ST_*. Estimates of *d_xy_* and nucleotide diversity were used to confirm genomic locations that were pinpointed as candidate selective sweep on the basis of values of Tajima’s *D* and *F_ST_*. Precisely, genomic regions were considered as targets of selection if they mapped to significant peaks or depressions of diversity and *d_xy_*, and their values of *F_ST_* and Tajima’s *D* were among the top 1% of the empirical distribution in at least one population. Significantly negative values of Tajima’s *D* relative to the genome-wide average suggest an increase in low-frequency mutations due to negative or positive selection whereas significantly positive values of Tajima’s *D* indicate a balancing selection.

**Figure 5.**
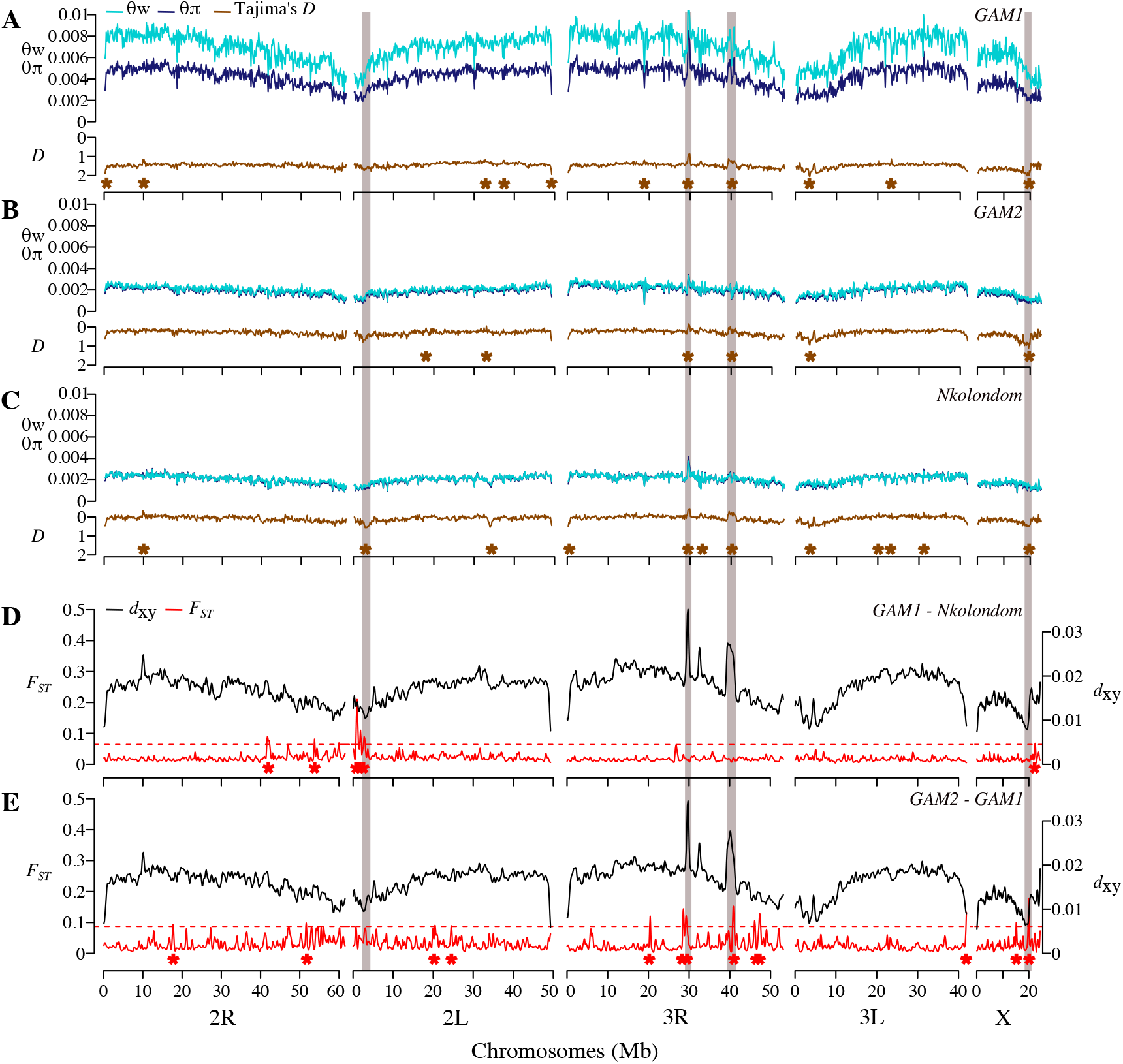
Genome scans reveal footprints of global and local adaptation in *An. gambiae* subpopulations. A-C) Diversity and Tajima’s *D* are plotted for each of the three subpopulations. Brown asterisks denote windows above the 99th percentile or below the 1st percentile of empirical distribution of Tajima’s *D*. D-E) Both absolute (*d_xy_*) and relative (*F_ST_*) divergence between populations are plotted across 150-kb windows. Red asterisks indicate windows above the 99th percentile of empirical distribution of *F_ST_*. In all populations, concordant dips in diversity and Tajima’s *D* are evident near the pericentromeric region of 2L where the *para* sodium channel gene is located. Three other selective sweeps located on the X and 3R chromosomes are highlighted (grey boxes).

**Figure 6.**
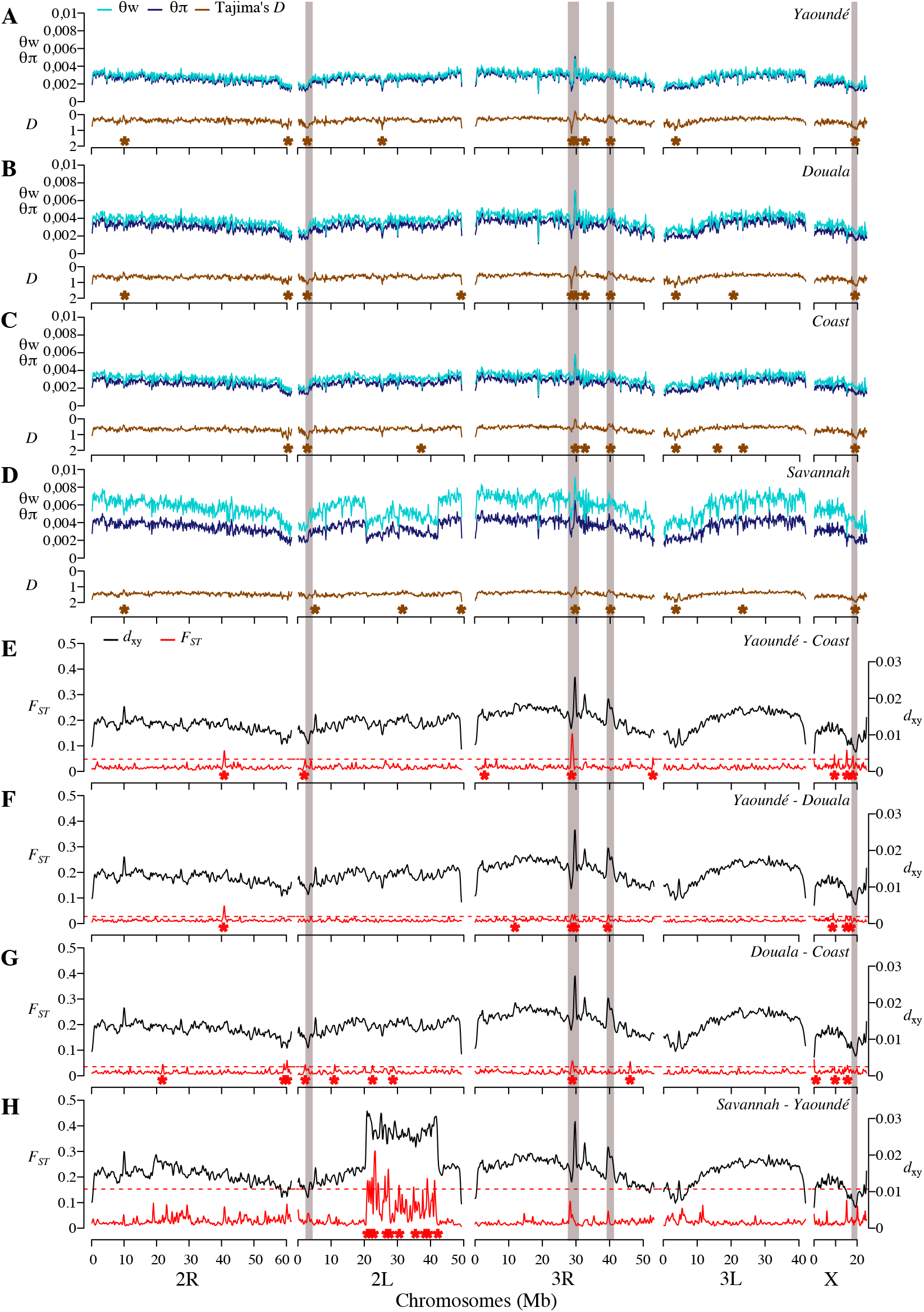
Strong signatures of selection are centered on xenobiotic resistance loci in subpopulations of *An. coluzzii*. A-H) Grey boxes highlight five selective sweeps characterized by extreme values of *F_ST_* and Tajima’s *D*. As in *An. gambiae*, sharp declines in diversity and allele frequency spectrum at the *para* sodium channel gene are present in all populations. A sweep encompassing a cluster of detoxification genes on 3R is sharpest in urban mosquitoes.

### *Targets of Selection within* An. gambiae *subgroups*

Our estimates of genome-wide diversity levels (Table S3) within *An. gambiae* subgroups are comparable to previous values based on RAD sequencing of East African *An. gambiae* s.l. populations (O’Loughlin et al. 2014).

As expected, the large *GAM1* population harbors more genetic diversity than the apparently rare *GAM2* population or the geographically restricted Nkolondom ecotype (Table S3, Figure 5A-C). *Tajima’s D* is consistently negative across the entire genome of all three subgroups, indicating an excess of low-frequency variants that are likely the result of recent population expansion (Figure 5A-C). Indeed, demographic models infer relatively recent bouts of population expansion in all three subgroups (Table S4).

Genome scans of each subgroup reveal genomic regions that show concordant reductions in diversity and allele frequency spectrum consistent with selection (highlighted in Figure 5 A-C). Based on the 1% cutoff of Tajima’s *D* (upper and lower bounds) and *F_ST_*, we identified 4 candidate regions exhibiting strong signatures of selection. It should be noted that due to the reduced representation sequencing approach we used, our analysis may be highlighting only clear instances of selection, which are likely both recent and strong (Tiffin and Ross-Ibarra 2014). An apparent selective event on the left arm of chromosome 2 near the centromere is found in all populations. This selective sweep is characterized by a prominent depression of Tajima’s *D* (~1.5Mb in width) and contains ~80 genes including the pyrethroid knockdown resistance gene (kdr). Although it is difficult to precisely identify the specific gene that has been influenced by selection in this region, the voltage-gated sodium channel gene (kdr), which is a pervasive target of selection in *Anopheles* mosquitoes (Clarkson et al. 2014; Norris et al. 2015) represents the strongest candidate in our populations. Three mutations of the para-type sodium channel conferring knockdown resistance (1014F, 1014L and 1014S) are widespread in Cameroon (Nwane et al. 2011). The drop in Tajima’s *D* in the putative *kdr* sweep is sharpest in the Nkolondom population. Both 1014F and 1014L *kdr* alleles coexist at high frequencies in mosquitoes from this suburban neighborhood of Yaoundé where larvae can be readily collected from irrigated garden plots that likely contain elevated levels of pesticides directed at agricultural pests (Nwane et al. 2011; Fossog Tene et al. 2013; Nwane et al. 2013).

Another region exhibiting consistent signatures of selection in all populations is found around the centromere on the X chromosome. Functional analyses of gene ontology (GO) terms (Table S5) revealed a significant representation of chitin binding proteins in this sweep (*p* = 1.91e-4). A strong genetic divergence among the three subgroups of *An. gambiae* characterized by significant *F_ST_* and *d_xy_* peaks is also observed at ~30 Mb and ~40 Mb on chromosome 3R. Positive outliers of both Tajima’s *D* and nucleotide diversity suggest that this genetic divergence is due to balancing selection on multiple alleles among populations. Functional analyses of GO terms indicate that the region at ~40 Mb on 3R is enriched in cell membrane proteins, genes involved in olfaction and epidermal growth factors (EGF) genes (Table S5). Finally, a striking depression in Tajima’s *D* supported by a marked dip in nucleotide diversity occurs on chromosome 2L from ~33-35 Mb in the Nkolondom population exclusively (Figure 5A-C). Despite the lack of genetic differentiation, this region – enriched in six EGFs (Table S5) – is probably a recent selective sweep, which could facilitate larval development of this subgroup in pesticide-laced agricultural water.

### *Targets of Selection within* An. coluzzii *subpopulations*

As above, we used diversity, allele frequency spectra and genetic differentiation metrics to scan for targets of selection in the four subgroups of *An. coluzzii*. Overall, genetic diversity is higher in the northern savannah population than either of three southern populations, which all exhibit similar levels of diversity (Table S3). Just as in *An. gambiae*, all subgroups have consistently negative Tajima’s *D* values confirming demographic models of population expansion (Table S4).

We detected 33 regions where either *F_ST_* or Tajima’s *D* was above the 99^th^ percentile of the empirical distribution in at least one population. This included at least 10 hotspots of *F_ST_* clustered within the 2L*a* inversion, which segregates between forest and savannah populations, as well as a region centered on the resistance to dieldrin (*rdl*) locus at ~25 Mb on 2L (Figure 6H). However, based on the 1% threshold of both Tajima’s *D* and *F_ST_*, we found only 5 candidate selective sweeps including the *kdr* region, the sweep on the X chromosome, and the two hot spots of balancing selection detected on the chromosome 3R in *An. gambiae* (Figure 6A-D). The fifth putative selective sweep characterized by a sharp drop in both diversity and Tajima’s *D* occurs on 3R from ~28.5-29.0 Mb, with the decline being more significant in urban populations. The particularly reduced diversity at this sweep in urban mosquitoes strongly suggests it may contain variant(s) that confer adaptation to human-induced changes. Indeed, this genomic region harbors a cluster of both Glutathione S-transferase (*GSTE1-GSTE7*) and cytochrome P450 (*CYP4C27, CYP4C35, CYP4C36*) genes, and functional analyses of GO terms reveals an overrepresentation of terms containing “Glutathione S-transferase” (Table S5, *p* = 5.14e-10). Both the *GSTE* and cytochrome P450 gene families are known to confer metabolic resistance to insecticides and pollutants in mosquitoes (Enayati et al. 2005; David et al. 2013). In particular, *GSTE5* and *GSTE6* are intriguing candidate targets of selection as each is up-regulated in highly insecticide resistant *An. arabiensis* populations that recently colonized urban areas of Bobo-Dioulasso, Burkina Faso (Jones et al. 2012).

Both relative and absolute divergences between populations at the two selective sweeps involved in xenobiotic resistance probably reflect a spatial variation in selection along gradients of anthropogenic disturbance. In particular, the *kdr* locus exhibits minimal divergence in all pairwise comparisons, suggesting that the same resistance haplotype is under selection in each population (Figure 6E-H). This hypothesis is corroborated by data on the distribution of *kdr* mutations in Cameroon (Nwane et al. 2011). As opposed to *An. gambiae* populations, which are broadly polymorphic for three *kdr* alleles, the 1014L mutation is the major allele present at very high frequencies in *An. coluzzii* throughout the country. The GSTE/CYP450 sweep on 3R shows a peak in *F*_ST_, but surprisingly low absolute divergence between Yaoundé and Coastal mosquitoes – a pattern consistent with genomic regions that underwent a selective sweep over a broad geographic area followed by episodes of within-population selection leading to reduced diversity across certain locations (Irwin et al. 2016). Comparisons between Douala and Coastal populations show a more moderate increase in *F*_ST_, presumably due to high rates of migration between these nearby sites. To further explore the 3R GSTE/CYP450 sweep, we reconstructed haplotypes for all 240 *An. coluzzii* southern chromosomes across the 28 SNPs found within the sweep. In the Yaoundé population, a single haplotype is present on 44 out of 80 (55%) chromosomes (all grey SNPs), while an additional 11 haplotypes are within one mutational step of this common haplotype (Figure 7A). In Douala, the same haplotype is the most common, but present at a lower frequency (31%) than in Yaoundé (Figure 7B). This haplotype is found on only 6/80 (7.5%) coastal chromosomes (Figure 7C). Population genomic analyses of the same 28 SNPs (Figure 7D-F) mirror results of the haplotype analysis, confirming that haplotype inference did not bias the results.

**Figure 7.**
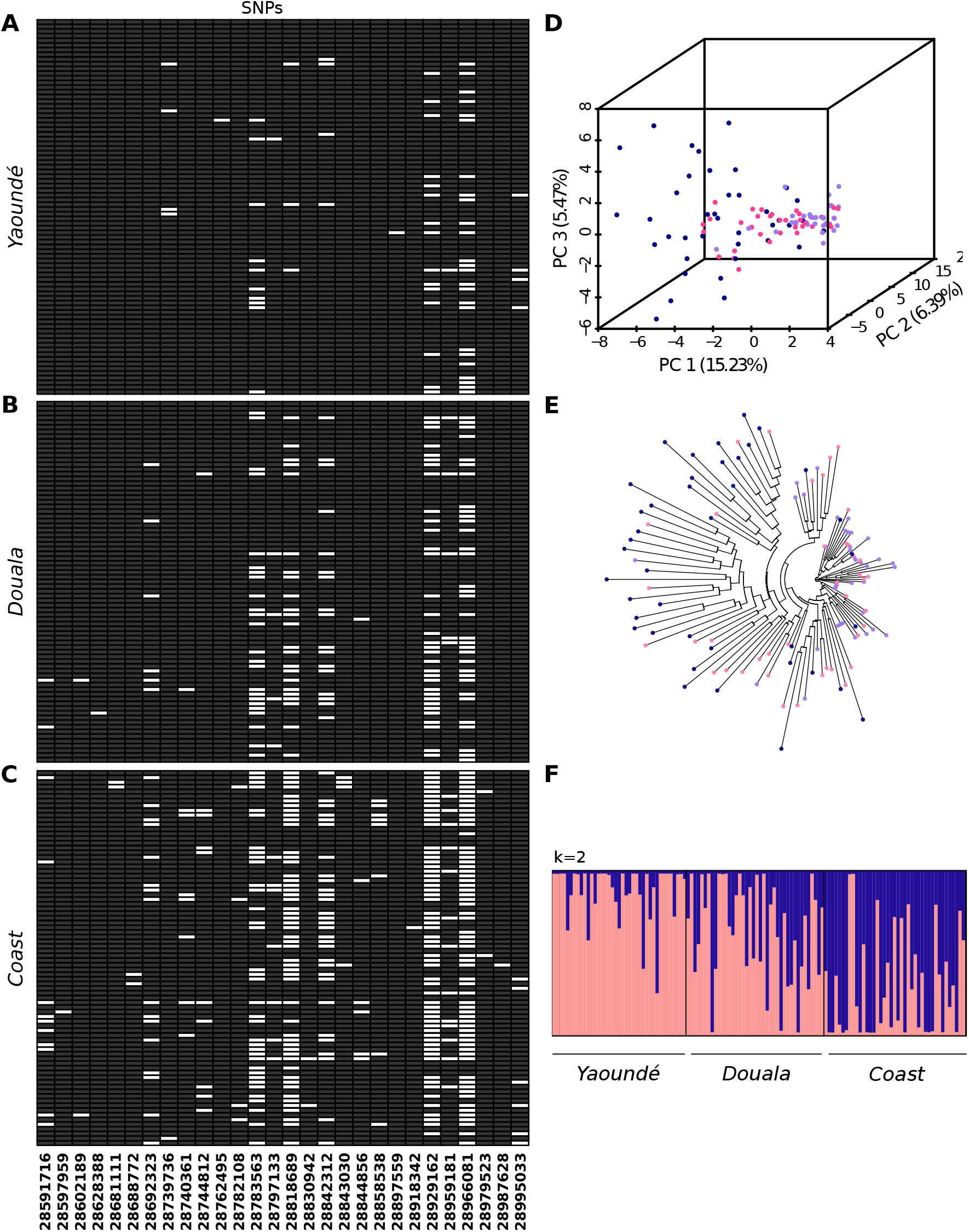
Spatially varying selection between urban and coastal populations. For each of the three southern *An. coluzzii* subpopulations, 80 reconstructed haplotypes are visualized by color-coding 28 bi-allelic SNPs in the 3R GSTE/CYP450 sweep either grey or white. A single invariant haplotype -- all grey SNPs -- is common in (A) Yaoundé, less so in (B) Douala, and very rare in (C) coastal populations. D-E) Similarly, in PCA and NJ Tree analysis of the same 28 SNPs, coastal individuals (navy blue) are diffuse across genotypic space, while Yaoundé mosquitoes (purple) are tightly clustered. As expected, Douala (pink) exhibits an intermediate degree of variation. F) STRUCTURE analysis based solely on the 28 SNPs within the sweep shows clear distinctions between the three populations.

## DISCUSSION

### *Genetic differentiation within species of the* An. gambiae *complex*

The *An. gambiae* complex is a model of adaptive radiation with a puzzling evolutionary history (Coluzzi et al. 2002; Ayala and Coluzzi 2005). Due to the lack of observable phenotypes that segregate between populations, it is almost impossible to tackle the genetic basis of divergence between and within species of the complex using traditional genetic mapping methods. Consequently, the examination of sequence divergence in the wild with high-throughput sequencing and genotyping approaches is particularly adapted to address the environmental and genomic targets of ecological divergence in *An. gambiae* s.l. In that respect, patterns of genomic divergence have started to be dissected and significant insights have been gained into the signatures of selection among continental populations (Lawniczak et al. 2010; Neafsey et al. 2010; White et al. 2011). However, ecological divergence and local adaptation within species of the *An. gambiae* complex sometimes occur at lower spatial scales (often a few kilometres) (Sogoba et al. 2008; Kamdem et al. 2012) that are ignored by continental-scale analyses. Here we have applied a population genomic approach to investigate the genomic architecture of selection across a more restricted geographic area characterized by a great environmental diversity. We first showed that reduced representation sequencing of 941 *An. gambiae* s.l. collected in or near human settlements in 33 sites scattered across Cameroon facilitated rapid identification of known sibling species and revealed a complex hierarchical population structure within *An. gambiae* and *An. coluzzii*. This result is opposite to that found in East Africa, where RADseq markers detected no population differentiation within species of the *An. gambiae* complex (O’Loughlin et al. 2014). Indeed, contrary to their East African counterparts, West African populations have a long history of genetic differentiation that is still unresolved despite the application of several generations of genetic markers (Coluzzi et al. 1979; Touré et al. 1998; Coluzzi et al. 2002; Wondji et al. 2002). Polymorphic chromosomal inversions and microsatellite markers have suggested the presence of additional subdivision within *An. coluzzii* in Cameroon (Wondji et al. 2005; Slotman et al. 2007). Using genome-wide SNPs, we have shown that this subdivision results in at least three genetically distinct clusters including populations from the savannah, coastal and urban areas. Little is known about the phenotypic differentiation of these populations, but recent studies suggest that mosquitoes from the coastal region tolerate higher concentrations of salt (Tene Fossog et al. 2015) while urban populations are highly resistant to insecticides and environmental pollutants (Antonio-Nkondjio et al. 2011; Fossog Tene et al. 2013; Tene Fossog et al. 2013; Antonio-Nkondjio et al. 2015). Models of ecological niches of *An. gambiae* s.l. in Cameroon predict that favourable habitats of *An. coluzzii* populations are fragmented and marginal in contrast to *An. gambiae*, which occupies a broad range across the country (Simard et al. 2009). In line with this prediction, we have found a weak differentiation and higher diversity in *An. gambiae* over significant geographic areas despite the presence of an emerging suburban cluster in Nkolondom.

### Genomic signatures of selection

When approached with powerful tools, sensitive enough to capture shallow and recent processes, populations depicting increasing levels of genetic differentiation along a speciation continuum may provide insights into the targets of selection at early stages of ecological divergence (Savolainen et al. 2013). In principle, weakly differentiated populations such as the subgroups we described typically display a marginal genomic divergence because there has not been enough time for neutral and selective forces to shape the genomic architecture (Nosil and Feder 2012; Andrew and Rieseberg 2013). Indeed, we found signatures consistent with selection only across a handful of loci in structured populations of *An. gambiae* and *An. coluzzii*. Although some targets of selection are likely missing due to the limited genomic coverage of the RADseq approach we used (Arnold et al. 2013; Tiffin and Ross-Ibarra 2014), the few selective sweeps we found are relevant based on the genes they contain. Specifically, the five main targets of selection in our populations are enriched in genes whose functions include insecticide resistance and detoxification, epidermal growth, cuticle formation and olfaction. The cuticle plays a major role at the interface between several biological functions in insects and recent data support the hypothesis that the coevolution of multiple mechanisms, including cuticular barriers, has occurred in highly pyrethroid-resistant *An. gambiae* (Balabanidou et al. 2016). Olfaction also mediates a wide range of both adult and larval behaviours including feeding, host preference, and mate selection in blood-feeding insects (Bowen 1991; Carey et al. 2010; Takken and Verhulst 2013).

Finally, due to the extreme association between the most effective vectors of the *An. gambiae* complex and human habitats, a long-standing hypothesis has suggested that human-driven selection is the main modulator of ecological divergence between and within species (Coluzzi et al. 1979; Coluzzi et al. 2002; Kamdem et al. 2012; Caputo et al. 2014). Consistent with this hypothesis, we have found that urban and suburban mosquitoes are genetically differentiated from rural populations and exhibit strong signatures of recent selection at loci containing xenobiotic resistance genes.

*Anopheles* mosquitoes currently experience strong selective pressures across the African continent due to the increased use of pesticides/insecticides in agriculture and vector control over the last few decades (WHO 2012; Reid and McKenzie 2016). The idea that insecticide resistance can affect the population structure of mosquitoes as a result of a limited gene flow and/or local adaptation between resistant and sensitive populations was already evoked in the 1990s (Silvestrini et al. 1998; Lenormand et al. 1999). In addition to increased resistance, large-scale exposure of vector populations to insecticide-treated bed nets in Africa is also correlated with changes in species distribution and behavioral shifts (Bøgh et al. 1998; Derua et al. 2012; Moiroux et al. 2012; Mwangangi et al. 2013; Sokhna et al. 2013). However, so far, there is no evidence that these changes can lead to genetic clustering or the creation of cryptic populations within species (Main et al. 2016). Our data also support that the genetic differentiation within the *An. gambiae* complex in Cameroon is due to regional-scale subdivisions that do not correspond to known distributional or behavioural shifts. Previous studies have documented significant reductions in the population size of *An. gambiae* s.l. after introduction of long lasting insecticide treated nets, but did not determine the influence of exposure on population or genomic structuring (Athrey et al. 2012). The demographic history of subgroups of *An. gambiae* and *An. coluzzii* inferred from allele frequency spectra does not indicate signals of recent population bottlenecks that can be associated with insecticide exposure (supplementary information, Table S4). It has also been shown that adaptive introgression, allowing genomic regions containing insecticide resistance genes to spread among sibling species of *An. gambiae* s.l., coincides with major insecticide-treated net distribution campaigns (Clarkson et al. 2014; Norris et al. 2015). Although the frequency of natural hybrids is low (less than 2%), episodic gene flow occurs between species of the complex in Cameroon (Lee et al. 2013), but adaptive introgression at insecticide resistance loci have yet to be explicitly demonstrated in this region. Insecticide resistance, involving both multiple alleles of *kdr* and significant levels of metabolic resistance, is widespread among *An. gambiae* and *An. coluzzii* populations in Cameroon especially in urban areas and locations where intensive farming practices are common (Ndjemaï et al. 2009; Antonio-Nkondjio et al. 2011; Nwane et al. 2011; Fossog Tene et al. 2013; Nwane et al. 2013; Antonio-Nkondjio et al. 2015). Alongside a mass distribution of insecticide-treated nets throughout the country that was held in 2011, indoor residual insecticide spraying has been tested in a few pilot locations, but this approach is not yet applied on a large scale (Etang et al. 2011; Bowen 2013). We hypothesized that regional variations in insecticide exposure exist across the country and contribute to the patterns observed at the two selective sweeps associated with xenobiotic resistance. Knockdown and metabolic resistance involving some of the genes present in the 3R GSTE/CYP450 sweep are the most common resistance mechanisms against the classes of insecticides currently used on a large scale that are found in *An. gambiae* s.l (Nwane et al. 2013). As expected, urban and suburban populations that are exposed to higher levels of insecticides/pollutants exhibit stronger signals of selection (Kamdem et al. 2012; Fossog Tene et al. 2013; Tene Fossog et al. 2013). Indeed, the synergistic effects of the two types of xenobiotics could be exerting intense selection pressure for pleiotropic resistance in urban mosquitoes (Nwane et al. 2013). Further analysis of the 3R GSTE/CYP450 sweep using a combination of whole genome resequencing and functional genomic approaches should help resolve the specific genes that are targets of local adaptation in this region of the genome. Additional studies are also needed to further characterize the spatial and temporal variations as well as the haplotypes correlated with insecticide-induced selection within the sweeps.

Effective strategies for managing insecticide resistance that has reached a critical tipping point among malaria vectors on the continent are urgently needed (Hemingway et al. 2016; Ranson and Lissenden 2016). In addition, considerable evidence suggests that insecticide resistance has much deeper consequences on the mosquito biology by interfering with other key biological functions such as the immune system, which in turn profoundly affects the fitness of the vector and its efficiency to transmit the pathogen (vectorial capacity) (Alout et al. 2013; Alout et al. 2014; Alout et al. 2016). As a result, detailed knowledge of genomic signatures of insecticide resistance and their pleiotropic effects on vectorial capacity is necessary to design effective resistance management. Surprisingly, despite intense research on insecticide resistance, this information is lacking in all malaria vectors partly because efforts have so far focused on a few resistance genes and have failed to capture the widespread effects of insecticide resistance across the genome. The population genomic approaches we have used are among the new tools that are being applied to investigate the genomic targets of insecticide resistance at a fine scale in mosquitoes and will facilitate the development of rational resistance management strategies.

## MATERIALS AND METHODS

### Mosquito collections

We collected *Anopheles* mosquitoes from 33 locations spread across the four major ecogeographic regions of Cameroon from August to November 2013 (Table S1). All collections were performed by researchers who were given malaria prophylaxis before and after the collection period. Indoor resting adult mosquitoes were collected by pyrethrum spray catch, while host-seeking adults were obtained via indoor/outdoor human-baited landing catch. Larvae were sampled using standard dipping procedures (Service 1993). Individual mosquitoes belonging to the *An. gambiae* complex were identified using morphological keys (Gillies and De Meillon 1968; Gillies and Coetzee 1987).

### ddRADseq Library Construction

Genomic DNA was extracted from adults using the ZR-96 Quick-gDNA kit (Zymo Research) and from larvae using the DNeasy Extraction kit (Qiagen). A subset of individuals were assigned to sibling species using PCR-RFLP assays that type fixed SNP differences in the rDNA (Fanello et al. 2002; Santolamazza et al. 2004). Preparation of ddRAD libraries largely followed (Turissini et al. 2014). Briefly, ~1/3^rd^ of the DNA extracted from an individual mosquito (10µl) was digested with *MluC1* and *NlaIII* (New England Biolabs). Barcoded adapters (1 of 48) were ligated to overhangs and 400 bp fragments were selected using 1.5% gels on a BluePippin (Sage Science). One of six indices was added during PCR amplification. Each library contained 288 individuals and was subjected to single end, 100 bp sequencing across one or two flow cells lanes run on an Illumina HiSeq2500.

Raw sequence reads were demultiplexed and quality filtered using the STACKS v 1.29 process_radtags pipeline (Catchen et al. 2011; Catchen et al. 2013). After removal of reads with ambiguous barcodes, incorrect restriction sites, and low sequencing quality (mean Phred < 33), GSNAP (Wu and Nacu 2010) was used to align reads to the *An. gambiae* PEST reference genome (AgamP3) allowing up to five mismatches per read. The STACKS pipeline was then used to identify unique RAD tags and construct consensus assemblies for each and to call individual SNP genotypes using a maximum-likelihood statistical model.

### Population Genomic Analysis

Population genetic structure was assessed using the SNP dataset output by the *populations* program of STACKS. We used PLINK v 1.19 to retrieve subsets of genome-wide SNPs as needed (Purcell et al. 2007). PCA, neighbor-joining tree analyses, and Bayesian information criterion (BIC) were implemented using the packages *adegenet* and *ape* in R (Paradis et al. 2004; Jombart 2008; R Development Core Team 2014). Ancestry analyses were conducted in fastSTRUCTURE v 1.0 (Raj et al. 2014) using the logistic method. The choosek.py script was used to find the appropriate number of populations (k); in cases where a range of k was suggested, the BIC-inferred number of clusters was chosen. CLUMPP v1.1.2 (Jakobsson and Rosenberg 2007) was used to summarize assignment results across independent runs and DISTRUCT v1.1 (Rosenberg 2004) was used to visualize ancestry assignment of individual mosquitoes. We used a subset of 1,000 randomly chosen SNPs to calculate average pairwise *F*_ST_ between populations in GENODIVE v 2.0 using up to 40 individuals – prioritized by coverage – per population (Meirmans and Van Tienderen 2004). Using this same subset of 1,000 SNPs, we conducted an AMOVA (Excoffier et al. 1992) to quantify the effect of the sampling method and the geographic origin on the genetic variance among individuals in GENODIVE. We used 10,000 permutations to assess significance of *F_ST_* values and AMOVA. We input pairwise *F_ST_* values into the program Fitch from the Phylip (Plotree and Plotgram 1989) suite to create the population-level NJ tree. F_*IS*_ values were computed with the *populations* program in STACKS.

### Genome Scans for Selection

We used ANGSD v 0.612 (Korneliussen et al. 2014) to calculate nucleotide diversity (*θ_w_* and *θ_π_*) and Tajima’s *D* in 150-kb non-overlapping windows. Unlike most genotyping algorithms, ANGSD does not perform hard SNP calls, instead taking genotyping uncertainty into account when calculating summary statistics. Similarly, absolute divergence (*d*_xy_) was calculated using *ngsTools* (Fumagalli et al. 2014) based on genotype likelihoods generated by ANGSD. Kernel smoothed values for 150-kb windows for all four metrics (*θ_w_, θ_π_*, *D*, *d*_xy_) were obtained with the R package *KernSmooth*. *F*_ST_ (based on AMOVA) was calculated with the *populations* program in STACKS using only loci present in 80% of individuals. A Kernel smoothing procedure implemented in STACKS was used to obtain *F*_ST_ values across 150-kb windows. Because regions with unusually high or low read depth can yield unreliable estimates of diversity and divergence parameters due to the likelihood of repeats and local misassembly, we checked that the average per-locus sequencing coverage was consistent throughout the genome (Figure S5). To determine if selective sweeps were enriched for specific functional annotation classes, we used the program DAVID 6.7 with default settings (Huang et al. 2009). We delimitated the physical limits of the selective sweep as the region corresponding to the base of the peak or the depression of Tajima’s *D*. Haplotypes across the GSTE/CYP450 sweep were reconstructed by PHASE v 2.1.1 using the default recombination model (Stephens et al. 2001; Stephens and Scheet 2005).

## ACKNOWLEDGEMENTS

This work was supported by the University of California Riverside and National Institutes of Health (1R01AI113248, 1R21AI115271 to BJW). We thank Elysée Nchoutpouem and Raymond Fokom for assistance collecting mosquitoes and Sina Hananian for assisting in DNA extraction. We thank the editor and three anonymous reviewers for their constructive comments, which helped us to improve the manuscript. This work would not have been possible without the collaboration of inhabitants and administrative authorities of all sampled sites and we want to underscore their generosity and patience.

## AUTHOR CONTRIBUTIONS

Conceived and designed the experiments: CK BJW. Performed the experiments: CK BJW SG. Analyzed the data: CK CF BJW. Wrote the paper: CK CF BJW.

## Supplemental Figure Legends

**Figure S1.**
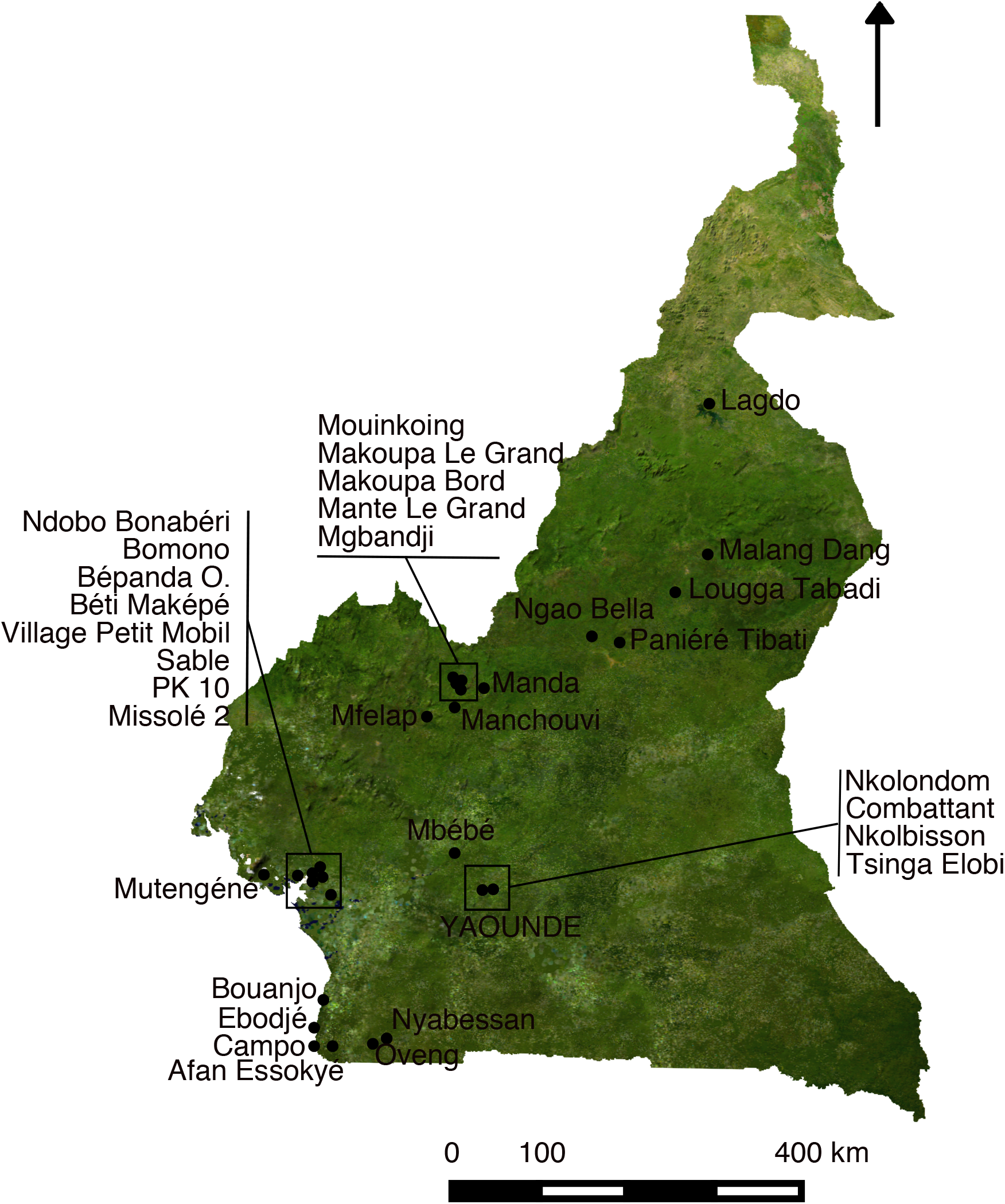
Detailed map of sampling sites in Cameroon.

**Figure S2.**
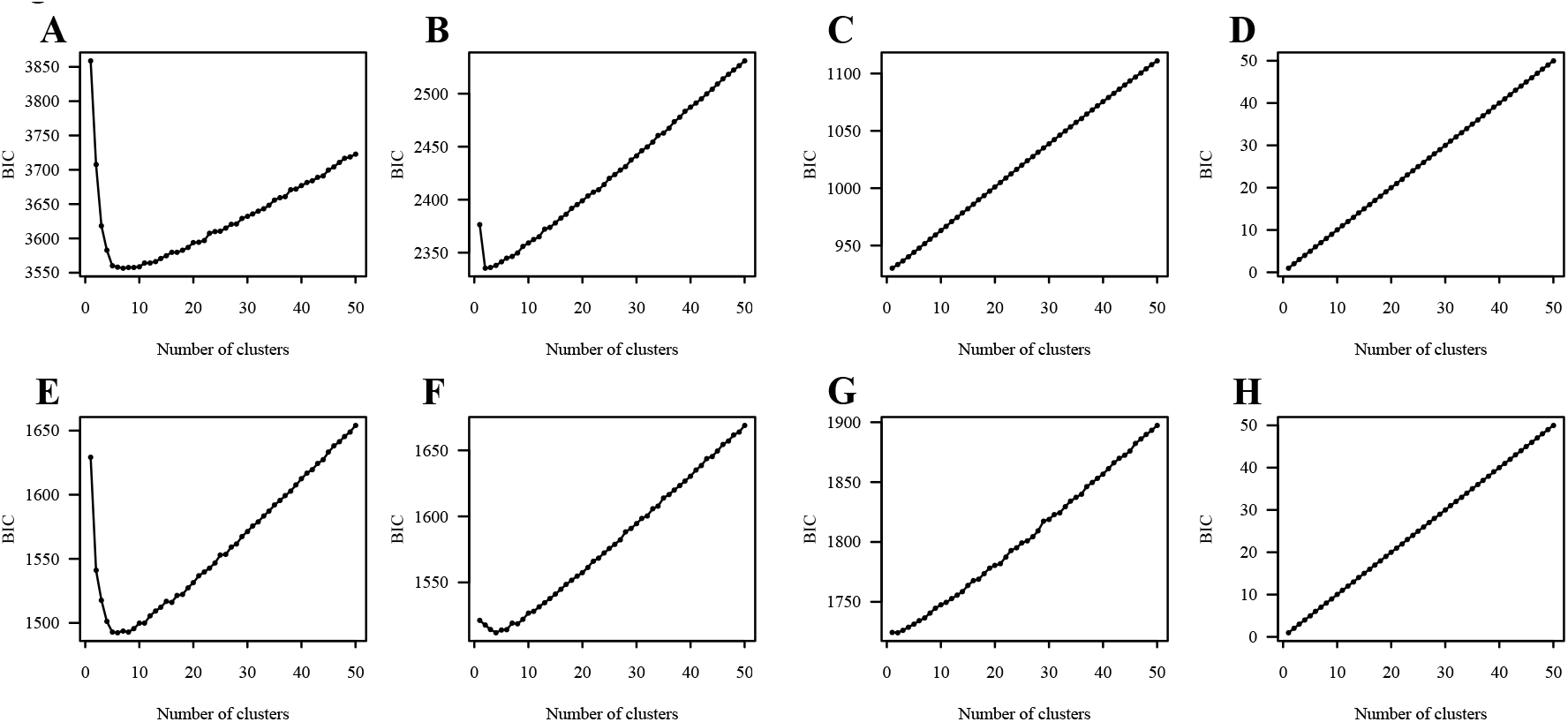
Bayesian information criterion was used to determine the most likely number of clusters/populations for A) all 941 samples, B) all *An. coluzzii*, C) *An. coluzzii* from the northern savannah, D) *An. arabiensis*, E) the 309 individuals used for genome scans F) all *An. gambiae*, G) southern *An. coluzzii*, and H) *An. melas*. BIC scores for 1 to 50 clusters are plotted. The lower the BIC score the better the model fits the observed genetic diversity.

**Figure S3.**
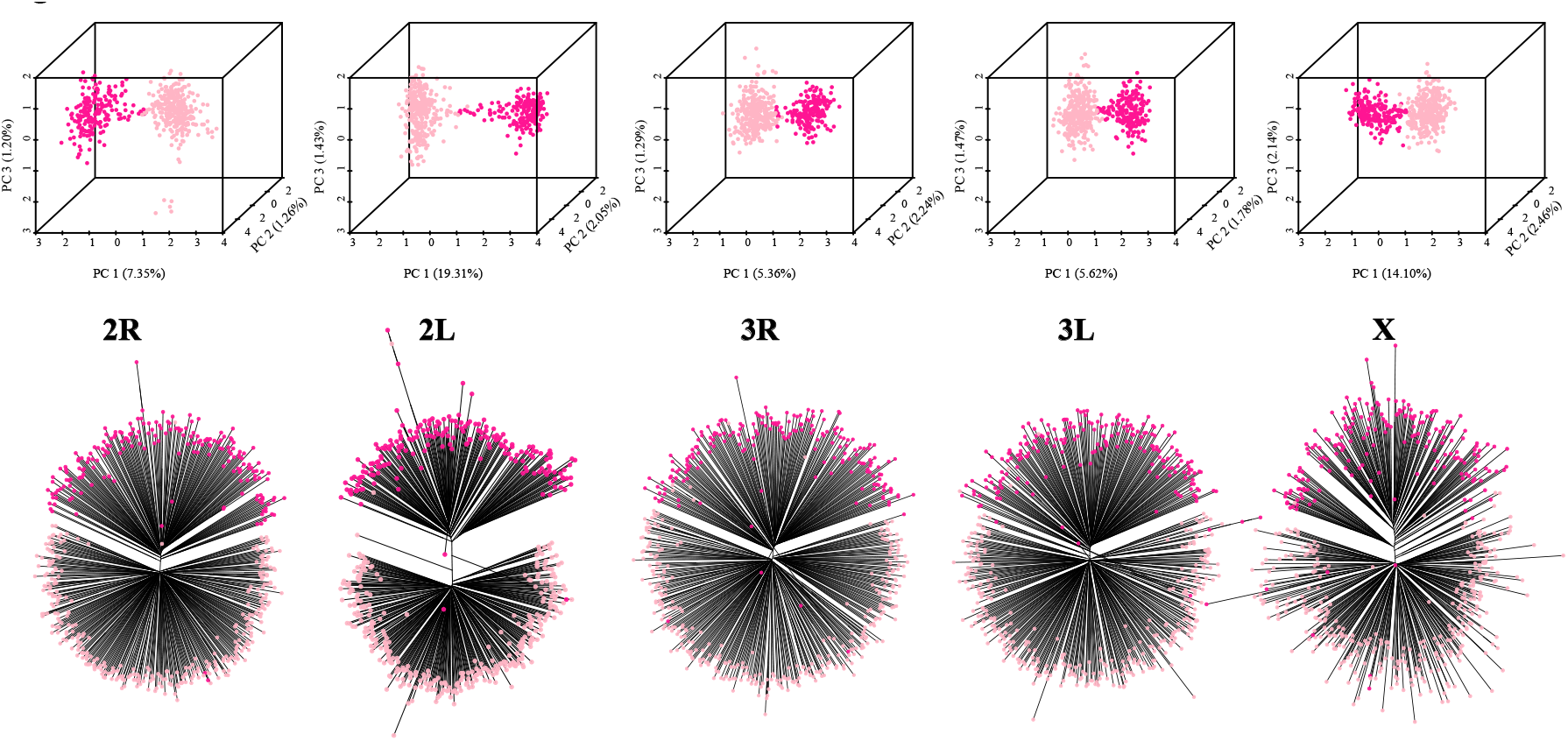
Southern (pale pink) and northern populations (deep pink) of *An. coluzzii* are readily separated in PCA (top) and NJ trees (bottom) using SNPs exclusively from any of the five chromosomal arms.

**Figure S4.**
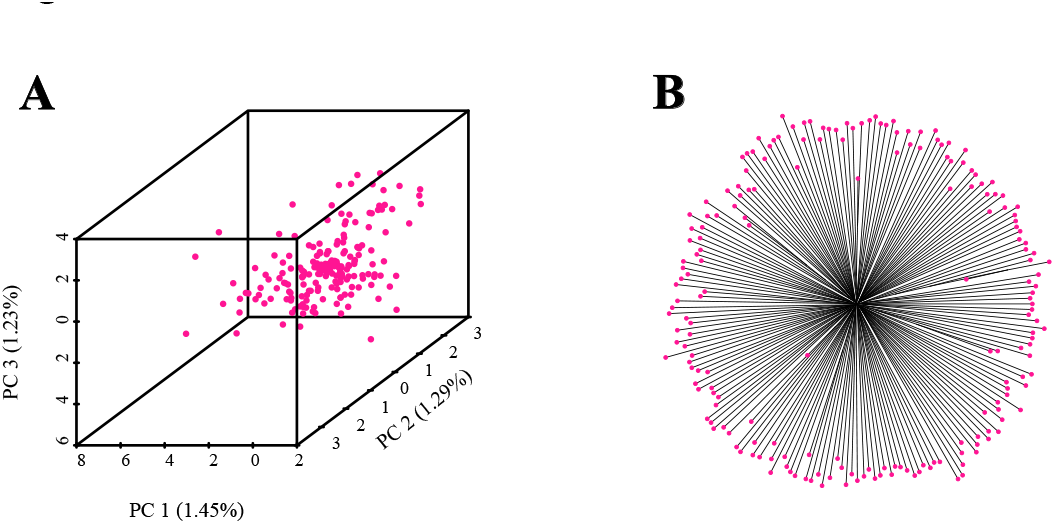
No population substructure is detectable within northern *An. coluzzii* using PCA and NJ tree analysis.

**Figure S5.**
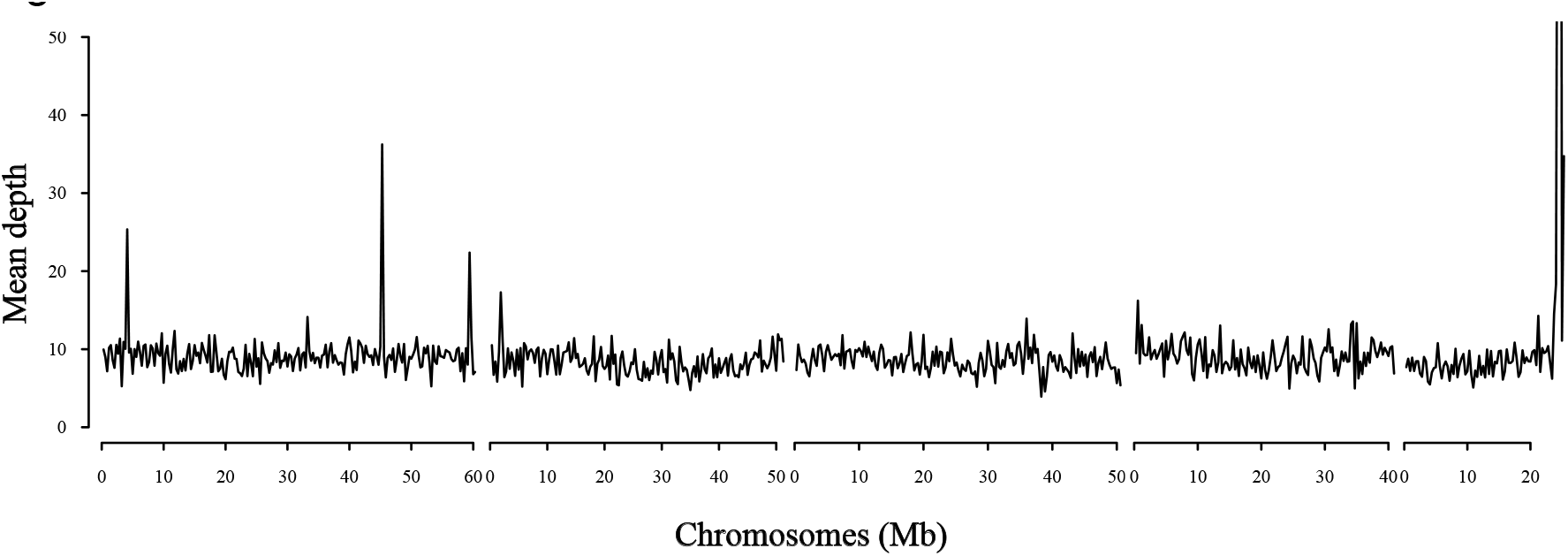
Mean sequencing coverage per individual is plotted in 300kb non-overlapping windows across the genome. Only individuals used in the genome scans are included in the coverage calculation.

## Supplemental Information

Consistently negative Tajima’s *D* across all subgroups may reflect recent population expansions. To further address this hypothesis we modeled the demographic history of each population using a diffusion-based approach implemented in the software package ∂a∂i v 1.6.3 (Gutenkunst et al. 2009). We fit four alternative demographic models (*neutral, growth, two-epoch, bottle-growth*), without migration or recombination, to the folded allele frequency spectrum of each cryptic subgroup of *An. gambiae s.s*. and *An. coluzzii*. The best model was selected based on the highest composite log likelihood, the lowest Akaike Information Criterion (AIC), and visual inspection of residuals. As the choice of model can be challenging in recently diverged populations, we prioritized the simplest model when we found it difficult to discriminate between conflicting models. To obtain uncertainty estimates for the demographic parameters we used the built-in bootstrap function implemented in dadi to derive 95% bootstrap confidence intervals.

Results indicate that GAM1, GAM2, and Savannah populations have experienced recent size increases. However, for the southern populations of *Yaoundé, Coast, Douala, and Nkolondom* the best demographic model is a *bottle-growth* (Table S4). While most classical studies report *An. gambiae s.l*. populations that are in expansion (Donnelly et al. 2001), a more recent study employing RAD markers revealed that some East African populations have more complex demographic histories, often involving several changes in effective population size (*Ne*) as we observed in southern forest populations of both *An. coluzzii* and *An. gambiae* (O’Loughlin et al. 2014).

---

**Table S1.**
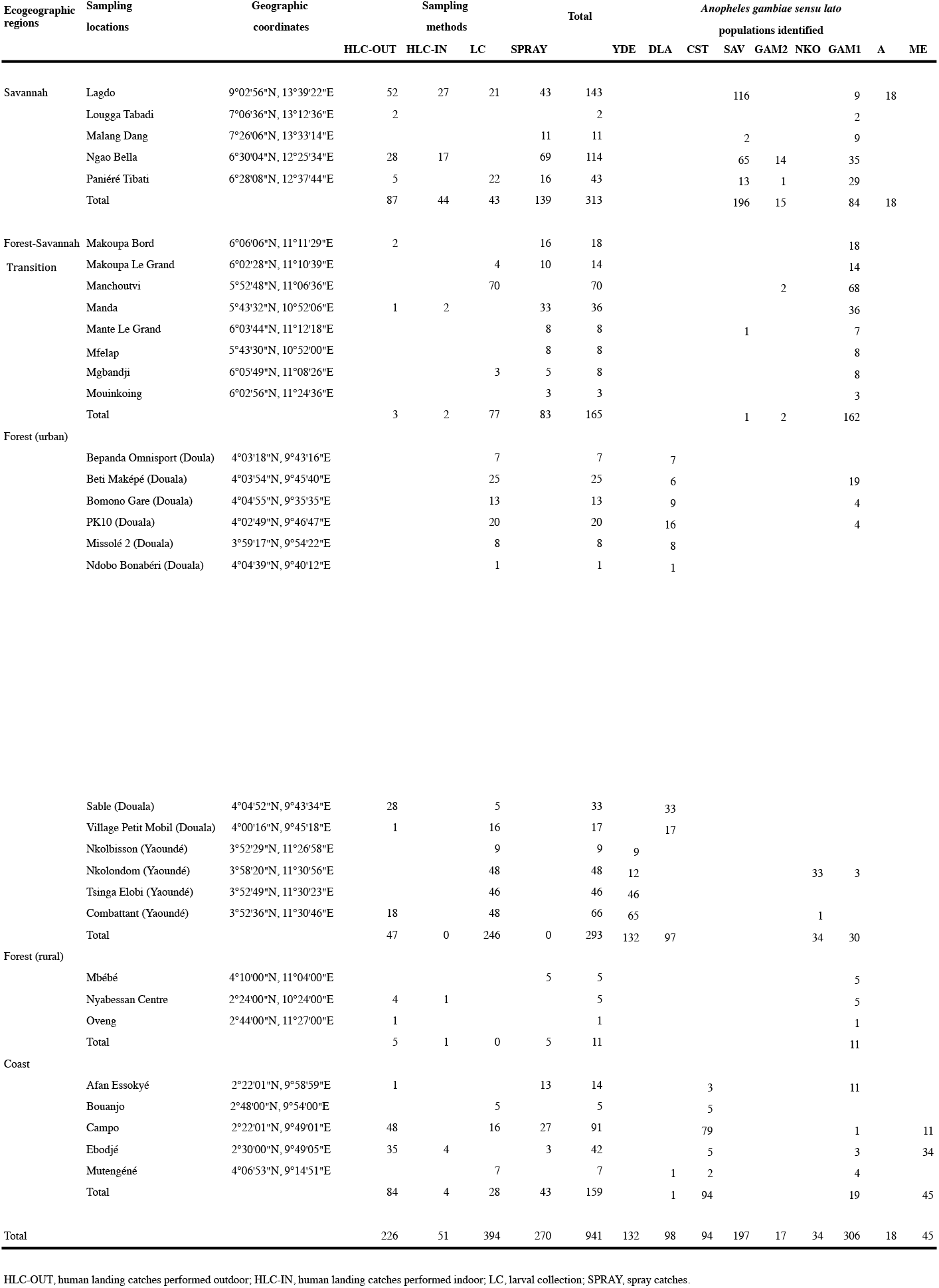
Locations of *An. gambiae s.l*. mosquitoes sequenced in Cameroon

**Table S2.** Pairwise comparison of genetic distance (*F*_ST_) among cryptic subgroups and sibling species of *An. gambiae s.l*.

| $F_{ST}$ | *Yaoundé* | *Douala* | *Coast* | *Savannah* | *GAM2* | *Nkolondom* | *GAM1* | *An. arabiensis* |
| --- | --- | --- | --- | --- | --- | --- | --- | --- |
| *Yaoundé* | - |  |  |  |  |  |  |  |
| *Douala* | 0.016 | - |  |  |  |  |  |  |
| *Coast* | 0.035 | 0.023 | - |  |  |  |  |  |
| *Savannah* | 0.127 | 0.109 | 0.103 | - |  |  |  |  |
| *GAM2* | 0.244 | 0.199 | 0.230 | 0.168 | - |  |  |  |
| *Nkolondom* | 0.197 | 0.201 | 0.200 | 0.247 | 0.161 | - |  |  |
| *GAM1* | 0.183 | 0.153 | 0.178 | 0.188 | 0.090 | 0.050 | - |  |
| *An. arabiensis* | 0.451 | 0.406 | 0.498 | 0.383 | 0.478 | 0.372 | 0.327 | - |
| *An. melas* | 0.851 | 0.836 | 0.830 | 0.792 | 0.841 | 0.818 | 0.781 | 0.872 |

**Table S3.** Average nucleotide diversity *gambiae s.s*.

| Population | $\theta\pi$ (bp<sup>-1</sup>) | $\theta w$ (bp<sup>-1</sup>) |
| --- | --- | --- |
| *Yaoundé* | 0.0025 | 0.0028 |
| *Douala* | 0.0031 | 0.0037 |
| *Coast* | 0.0025 | 0.0031 |
| *Savannah* | 0.0035 | 0.0057 |
| *GAM2* | 0.0019 | 0.0020 |
| *Nkolondom* | 0.0020 | 0.0021 |
| *GAM1* | 0.0042 | 0.0069 |

**Table S4.**
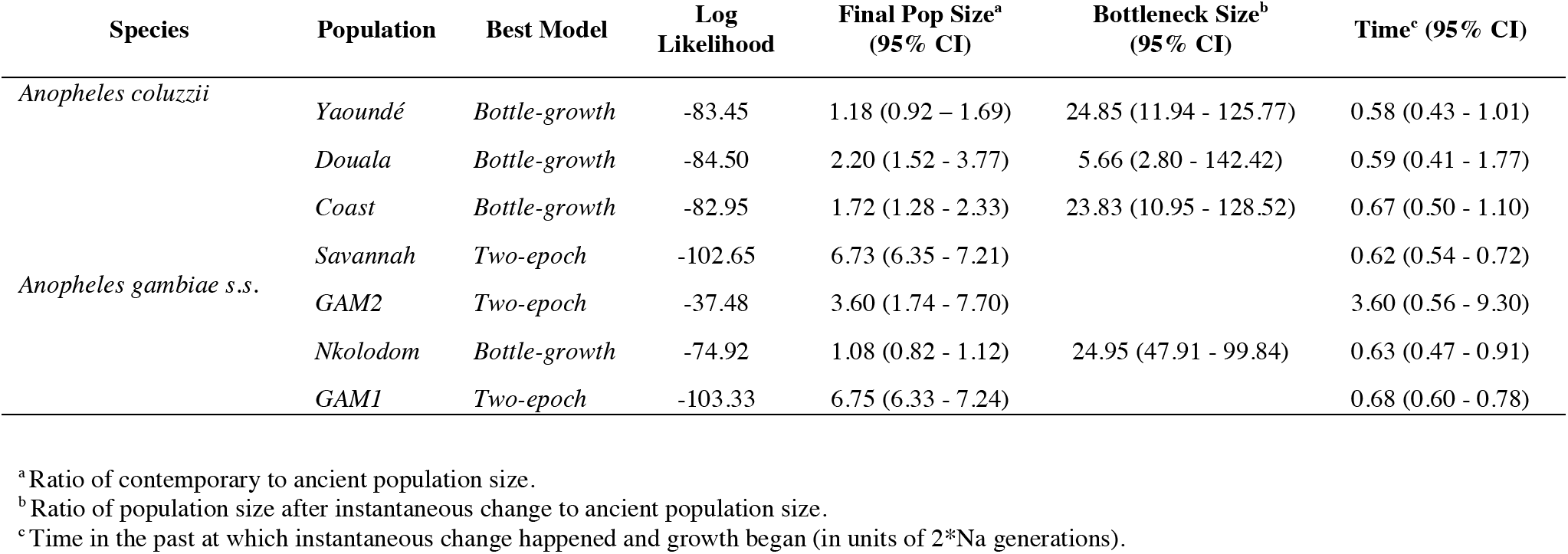
Parameters of demographic models inferred from folded Site Frequency Spectrum (SFS) of autosomal SNPs in seven cryptic subpopulations of *An. gambiae*

**Table S5.** Functional analysis of gene ontology terms in candidate regions. Supplementary online information

